# Structural brain connectivity in children with developmental dyscalculia

**DOI:** 10.1101/2021.04.16.440038

**Authors:** Nazife Ayyıldız, Frauke Beyer, Sertaç Üstün, Emre H. Kale, Öykü Mançe Çalışır, Pınar Uran, Özgür Öner, Sinan Olkun, Alfred Anwander, A. Veronica Witte, Arno Villringer, Metehan Çiçek

## Abstract

Developmental dyscalculia (DD) is a neurodevelopmental disorder specific to arithmetic learning even with normal intelligence and age-appropriate education. Difficulties often persist from childhood through adulthood. Underlying neurobiological mechanisms of DD, however, are poorly understood. This study aimed to identify possible structural connectivity alterations in DD. We evaluated 10 children with *pure* DD (11.3 ± 0.7 years) and 16 typically developing (TD) peers (11.2 ± 0.6 years) using diffusion tensor imaging. We first assessed white matter microstructure with tract-based spatial statistics. Then we used probabilistic tractography to evaluate tract lengths and probabilistic connectivity maps in specific regions. At whole brain level, we found no significant microstructural differences in white matter between children with DD and TD peers. Also, seed-based connectivity probabilities did not differ between groups. However, we did find significant differences in regions-of-interest tracts which had previously been related to math ability in children. The major findings of our study were reduced white matter coherence and shorter tract lengths of the left superior longitudinal/arcuate fasciculus and left anterior thalamic radiation in the DD group. Furthermore, lower white matter coherence and shorter pathways corresponded with the lower math performance as a result of the correlation analyses. These results from regional analyses indicate that learning, memory and language-related pathways in the left hemisphere might underlie DD.

## 1. INTRODUCTION

Developmental dyscalculia (DD or mathematical learning disability) is a neurodevelopmental disorder that negatively affects learning of numerical concepts and operations despite normal intellectual functioning (Adler, 2001; American Psychiatric Association, 2013; Butterworth, 2004; Shalev, 2004). Prevalence of pure DD has been estimated to be 3–8% of the general population, depending on the identification criteria, and this rate increases up to 13.8% when comorbidities (e.g., attention deficit hyperactivity disorder (ADHD), dyslexia) are present (Barbaresi et al., 2005; Butterworth & Laurillard, 2010; Shalev & von Aster, 2008). Clinical features of dyscalculia include use of “primitive” strategies to solve math problems with high error rates and long delays, and poor performance in symbolic and non-symbolic magnitude comparison tasks (Geary, 1993; Landerl et al., 2004; Schwenk et al., 2017; Shalev & Gross-Tsur, 2001). Educational achievements in children with DD are typically far behind their peers, and difficulties often continue into adulthood by negatively affecting academic career and quality of life (Gerber, 2012; Wilson et al., 2015). Despite its importance, the origins and biological basis of pure DD have not yet been clarified.

Numerical abilities have been localized to fronto-parietal and occipito-temporal networks including hippocampus, insula and claustrum in the human brain (Arsalidou et al., 2018; Arsalidou & Taylor, 2011; Houde et al., 2010; Moeller et al., 2015; Peters & De Smedt, 2018; Sokolowski et al., 2017). Alterations in brain connectivity patterns between these regions have been suggested to be related to DD (Arsalidou et al., 2018; Kaufmann et al., 2011; Matejko & Ansari, 2015).

A widely used *in-vivo* approach to study such alterations of structural connectivity is diffusion tensor imaging (DTI). The method assesses the diffusivity of water which is higher parallel than perpendicular to nerve fibers (Le Bihan et al., 2001; Mori & Barker, 1999). Therefore, DTI provides information about white matter (Buchsbaum et al.) microstructure (i.e., fractional anisotropy (FA), axial, radial and mean diffusivities (AD, RD, MD)) and it is possible to model the brain fibers via tractography (Basser, 1995; Feldman et al., 2010).

So far, two DTI studies have compared brain structural connectivity between children with DD and typically developing (TD) peers. Rykhlevskaia et al. (2009) found that 7–9-year-old children with DD showed reduced FA in the right corticospinal tract (CST), right inferior longitudinal fasciculus (ILF) and right inferior fronto-occipital fasciculus (IFOF), bilateral anterior thalamic radiation, bilateral superior longitudinal fasciculus (SLF) and forceps major and splenium of the corpus callosum (CC), regions which also showed reduced WM volumes compared to TD children. In the same study, with probabilistic and deterministic tractography, TD children showed stronger connectivity in interhemispheric superior parietal connections and right fusiform gyrus connections with the right temporo-parietal cortex (Rykhlevskaia et al., 2009). The second study reported reduced FA and increased RD in the posterior part of the left SLF in a voxel-wise whole brain as well as regions-of-interest based analysis in 10-year-old children with DD compared to TD peers (Kucian et al., 2014).

Other studies have provided evidence for the role of association fibers in numerical abilities of TD children. Tsang et al. (2009) found that FA values in the central part of the left anterior SLF were positively correlated with performance in the approximate arithmetic task of 10–15-year-old children. Additionally, they showed positive FA correlations in the SLF, ILF and IFOF bilaterally in tract-based spatial statistics (TBSS) whole brain analysis with arithmetic performance. Positive correlations between FA values in the left ILF and performances in numerical operations were detected in 7–9-year-old children (Van Eimeren et al., 2008). Fiber tracking analysis with the left intra parietal sulcus revealed connections with the left SLF, bilateral ILF and IFOF, and FA values in these regions were positively correlated with arithmetic scores in 9.6–11.3-year-old children (Li, Hu, et al., 2013). A whole brain tractography study showing higher FA values in the left anterior arcuate fasciculus (AF) specifically predicted better addition and multiplication in 11.5–12.9-year-old children (Van Beek et al., 2014).

In addition to association fibers, projection and commissural connections are related to mathematical competencies in TD children. FA values in the left superior corona radiata (SCR) were positively correlated with performance in numerical operations and mathematical reasoning tasks (Van Eimeren et al., 2008). FA values in the internal and external capsules bilaterally (IC and EC) were found to be positively correlated with performance in approximate addition (Tsang et al., 2009). In 6-year-old children, higher FA values in the left isthmus (body) of the CC, namely, interhemispheric IPS tracked via deterministic tractography, predicted higher performance in the numerical comparison tasks (Cantlon et al., 2011). Li, Hu, et al. (2013) and Tsang et al. (2009) showed that children’s arithmetic performances were positively correlated with FA values in the forceps major and the splenium of the CC, respectively.

So far, the relationship of tract length with numerical abilities has not been studied. Some developmental and aging studies have shown that tract length tends to become longer from birth to young adulthood and shorter again during normal aging (Marner et al., 2003; Tang et al., 1997; Yu et al., 2014). Several studies showed that neuropsychiatric diseases correlate negatively with tract length (Buchsbaum et al., 2006; Heaps-Woodruff et al., 2018; Salminen et al., 2013; Torgerson et al., 2013). It seems plausible to hypothesize that impaired numerical abilities of dyscalculic children correspond with shorter tract lengths as compared to their peers.

Taken together, previous studies indicate that the WM coherence of association (e.g., SLF, ILF, IFOF), projection (e.g., CST, ATR, SCR) and commissural (e.g., parts of CC) fibers correlate with numerical abilities in children (Matejko & Ansari, 2015; Moeller et al., 2015; Peters & De Smedt, 2018). To provide more evidence for the hypothesis that DD corresponds with altered structural connectivity, in this study we measured WM microstructure properties, lengths of the reconstructed tracts and connection probability via probabilistic tractography in children with pure DD and TD peers.

## 2. METHODS

### 2.1. Participants and Procedure

Participants were recruited in three stages from 13 primary state schools of low, medium and high socioeconomic backgrounds in Ankara, the capital of Turkey. In the first stage, we screened 2058 third-grade students from these schools with fluid intelligence (Raven’s Standard Progressive Matrices Test (RSPM) (Raven, 2000); Turkish version by Şahin and Düzen (1993)) and mathematics ability tests (curriculum based Mathematics Achievement Test (MAT) (Fidan, 2013) and Calculation Performance Test (CPT) (De Vos, 1992); Turkish version by Olkun et al. (2013)). The children’s teachers completed ADHD (attention deficit hyperactivity disorder) and learning difficulty questionnaires (Swanson, Nolan, and Pelham, SNAP-IV Questionnaire; reliability and validity by Bussing et al. (2008) and Strengths and Difficulties Questionnaire (SDQ) by Goodman (2001); Turkish version by Güvenir et al. (2008)). In the second stage, children at risk of developmental dyscalculia (DD candidates) and children with normal achievement as typically developing (TD candidates) were determined via inclusion and exclusion criteria according to the results of the screening tests and questionnaires. First, all children with incomplete data or those outside the mean age range of ±2.5 SD were excluded, leaving 1880 possible participants (mean age = 8.66±0.41, 953F, 927M). We divided the age range (7.5–9.5 years) into four categories of six-month periods. From this participant pool we determined 235 DD candidates and subsequently 235 TD candidates by applying inclusion and exclusion criteria within each six-month age category: DD candidates were those in the lowest 25th percentile of the MAT and CPT scores; TD candidates were chosen randomly between 35th and 75th percentiles of the MAT and CPT scores. Exclusion criteria for all participants included: 1) the lowest 10th percentile of the RSPM scores, and 2) the highest 15th percentile of the SNAP-IV scores.

In the last stage, two years later, we re-assessed DD and TD candidates in detail. We excluded children who were left-handed, born prematurely (birth weight < 2500g and gestation period < 36w) or those with psychiatric or neurological diseases. Child and adolescent psychiatrists evaluated volunteer participants to determine psychiatric comorbidities, such as ADHD and anxiety, during semi-structured interviews (Schedule for Affective Disorders and Schizophrenia for School Age Children-Present and Lifetime Version-Turkish Version) (Gökler et al., 2004). Expert psychologist tested the children using the Wechsler Intelligence Scale for Children (WISC-R) (Wechsler, 1974); Turkish version by Savaşır and Şahin (1995)) to exclude those with mental retardation (total IQ score < 80). We re-tested the children with the MAT and CPT to ensure they were in the previously determined groups. We evaluated reading disability with a time-limited reading test and eliminated dyscalculic children who read less than 80 words per minute (Öner et al., 2019). All of the detailed evaluations led to the exclusion of 12 participants; 14 participants did not agree to the MRI assessment and we were not able to reach all of the candidates in the determined pool. Therefore, we included 12 DD and (8F, mean age=11.2±0.7 years old) and 16 TD (9F, mean age=11.2±0.6 years old) children in the MRI part of the study. We additionally had to exclude two DTI data sets from the DD group as explained in Section 2.3. Demographic and cognitive profiles of the sample can be seen in Table 1. DD children had significantly lower MAT and CPT scores than TD peers, whereas age, gender, handedness, total WISC-R (corrected for arithmetic) and reading scores were well-matched between the groups.

**Table 1.** Demographics and behavioral cognitive profiles of the developmental dyscalculia (DD) and typically developing (TD) groups.

|  | **DD** | **TD** |
| --- | --- | --- |
| N | 10 | 16 |
| Gender (f/m) | 8/2<sup>a, e</sup> | 9/7<sup>a, e</sup> |
| Handedness (r/l) | 10/0 | 16/0 |
| Handedness | 13.8 (1.1)<sup>b, e</sup> | 14.5 (1.8)<sup>b, e</sup> |
| Age (years) | 11.3 (0.7)<sup>b, e</sup> | 11.2 (0.6)<sup>b, e</sup> |
| Age range (years) | 10.1 – 12.2 | 10.4 – 12.2 |
| Numerical Tests |  |  |
| MAT | 9.1 (2.9)<sup>**b</sup> | 20.3 (2.5)<sup>**b</sup> |
| CPT Sum | 67 (16.3)<sup>**b</sup> | 104.6 (10.7)<sup>**b</sup> |
| CPT Addition | 19 (3.8)<sup>**c</sup> | 25.4 (3.3)<sup>**c</sup> |
| CPT Subtraction | 14.9 (4.7)<sup>*b</sup> | 20.4 (2.8)<sup>*b</sup> |
| CPT Multiplication | 13.2 (5.1)<sup>**c</sup> | 21.6 (2.7)<sup>**c</sup> |
| CPT Division | 7.4 (3.8) <sup>**b</sup> | 16.6 (2.8) <sup>**b</sup> |
| CPT Mix | 12.5 (3) <sup>**b</sup> | 20.5 (3.1) <sup>**b</sup> |
| IQ Tests |  |  |
| WISC-R Performance | 98.6 (9.7)<sup>b, e</sup> | 103.3 (11)<sup>b, e</sup> |
| WISC-R Verbal | 88.3 (9) <sup>**b</sup> | 106.6 (10.5) <sup>**b</sup> |
| Arithmetic Sub Test (from Verbal IQ) | 6.5 (2.1)<sup>*b</sup> | 10.6 (2.9)<sup>*b</sup> |
| WISC-R Total | 92.5 (8.1)<sup>*b</sup> | 105.7 (9.6)<sup>*b</sup> |
| WISC-R Total (corrected for Arithmetic) | 95.6 (SE=3.2)<sup>d, e</sup> | 103.8 (SE=2.4)<sup>d, e</sup> |
| Reading Evaluation | 101.6 (12.9)<sup>c, e</sup> | 106.7 (25.7)<sup>c, e</sup> |

MAT: Mathematics Achievement Test, CPT: Calculation Performance Test, WISC-R: Wechsler Intelligence Scale for Children, Handedness Scale (Nalçacı et al., 2002; adapted from Chapman and Chapman, 1987). Scores are given as mean where appropriate. <sup>a</sup>: Fischer-Chi-Square-test, <sup>b</sup>: Independent-samples t-test, <sup>c</sup>: Mann-Whitney-U-test, <sup>d</sup>: ANCOVA. \*: p <0.05, \*\*: p<0.001, <sup>e</sup>: p>0.05.

Behavioral tests (MAT, CPT, RPM, SNAP IV and SDQ Questionnaire) used in this study were described in detail by Öner et al. (2018).

### 2.2. DTI and T1-weighted data acquisitions

Participants were familiarized with the MRI procedure in a mock scanner. We acquired anatomical and diffusion MRI data on a 3 Tesla Siemens Magnetom Tim Trio MRI scanner (Siemens Healthcare, Erlangen, Germany; software syngo MR B17) using a 16 channel head-coil. Children watched cartoons via the projective mirror system in the MRI scanner during both acquisitions. High-resolution diffusion-weighted images were obtained along 60 different directions with a b-value of b=700s/mm^2^ and ten initial reference images with b=0s/mm^2^. DTI data were acquired with anterior-to-posterior phase encoding direction containing 75 slices in transversal orientation without gap. An echo-planar imaging sequence was used via interleaved recording. Other scan parameters were: TR: 9.506s, TE: 85ms, FOV: 198mm, Flip Angle: 90, voxel size 1.55mm x 1.55mm, slice thickness 1.6mm, single spin echo. GRAPPA method with acceleration factor 2 and fast Gradient mode was used. DTI acquisition lasted 11m 35s. High-resolution structural T1-weighted image acquisition parameters were: TR: 2.6s, TE: 3.02ms, FOV: 256 mm, matrix: 256×256 and slice thickness: 1mm.

### 2.3. DTI Data Pre-processing

Pre-processing was implemented in a Nipype workflow (Gorgolewski et al., 2011). Firstly, DTI data were corrected with the MRtrix3 (https://www.mrtrix.org, (Tournier et al., 2019) DWI-denoising tool to improve SNR and estimation of diffusion parameters and fiber orientations by suppressing local signal fluctuations due to thermal noise (Veraart, Fieremans, et al., 2016; Veraart, Novikov, et al., 2016). Then we corrected the data for eddy-current induced distortions and subject head motion with the software “eddy” (Andersson & Sotiropoulos, 2016) implemented in FSL (https://fsl.fmrib.ox.ac.uk/fsl v6.0.1, (Smith et al., 2004). Simultaneously, we performed slice-to-volume correction (Andersson et al., 2017), and outlier detection and replacement (Andersson et al., 2016). Because children, especially those with developmental difficulties (Dosenbach et al., 2017; Yendiki et al., 2014) tend to move more during MRI, we used stringent criteria in these corrections.

We performed automated quality control on the single subject (QUAD) and group level (SQUAD) to detect data acquisition and pre-processing issues based on the output of the “eddy” tool (Bastiani et al., 2019). According to these results, we decided to exclude two data sets which had the highest motion parameters and outlier replacement percentage as well as the lowest SNR/CNR (see Supplementary Material, Table S1 and Fig S1). Using motion parameters as nuisance regressor in analysis or balancing them between groups reduces confounding effects of motion between groups (Yendiki et al., 2014). We compared absolute and relative motion parameters between DD and TD groups with independent samples t-test in SPSS software v24. The exclusion of the two datasets led to more balanced groups with respect to subject motion (for t-test results and detailed information see Supplementary Material, Table S1, Notes S1 and S2).

After eddy-current and motion correction, we computed a brain mask from the first non-diffusion weighted (b0) images using the Brain Extraction Tool (BET) form FSL (fractional intensity threshold = 0.2, with iteration method) (Smith, 2002). Using FSL, we calculated the tensor model at each voxel from the diffusion-weighted images and obtained Fractional Anisotropy (FA), Axial Diffusivity (AD), Radial Diffusivity (RD) and Mean Diffusivity (MD) maps. FA represents the degree of anisotropic diffusion in tissue (Alexander, Lee, Lazar, & Field, 2007). While MD measures the average diffusivity in tissue, AD represents the diffusivity along main fiber direction and RD describes the average diffusivity perpendicular to the axonal axis. These DTI metrics are commonly used to characterize WM microstructure. They are influenced by different microstructural properties such as axonal density, white matter coherence and/or maturation, the intra voxel orientation dispersion, myelination, membrane permeability or the number of axons (Feldman et al., 2010; Jones et al., 2013). For example, tracts with a denser axonal packing and higher myelinations have higher FA and lower RD compared to less densely packed tracts (Feldman et al., 2010). Higher FA and/or lower diffusivities are therefore often interpreted as higher WM coherence relating to faster and more efficient transmission of the axonal signals.

### 2.4. DTI Data Analyses

#### 2.4.1. Voxel-wise whole brain analysis in Tract-Based Spatial Statistics (TBSS)

We inspected WM microstructure with individual FA, MD, RD and AD maps performing voxel-wise whole brain analysis with TBSS (Smith et al., 2006). Because there have been only two dyscalculia studies with different results, samples and analysis methods, we first analyzed possible group differences on the whole brain level. We performed TBSS with its standard steps. First, we computed non-linear registration between all FA images and chose the structurally most representative FA map among all subjects as target image as suggested in TBSS for child data. We then aligned all FA images to the MNI152 1×1×1mm standard space by computing an affine registration of the target image to MNI and by applying the two combined registration steps for all images. Afterwards, all images were averaged and the skeleton of the mean FA image was created. The appropriate threshold for the mean FA skeleton was determined as 0.2 for the mean FA skeleton mask. Finally, all FA and non-FA data (MD, RD and AD) were projected onto the mean FA skeleton and the individual skeletonized FA, MD, RD and AD maps were obtained for further voxel-wise analyses.

The FSL tool “randomise” was used with permutation test for the TBSS voxel-wise analyses as suggested for GLM models (Winkler et al., 2014). Threshold-Free Cluster Enhancement (TFCE) method was used for statistical analysis (Smith & Nichols, 2009). Ten thousand permutations were performed for each independent samples t-test. Statistical significance value was defined as p<0.05 and Family-Wise Error correction (FWE) for multiple comparisons was applied in this p-value. The contrast of Controls>Dyscalculics demonstrated which WM tracts showed higher FA, MD, RD and AD values in TD controls than children with DD. The contrast of Dyscalculics>Controls demonstrated vice versa.

#### 2.4.2. Region of Interest (ROI) Analyses in TBSS

We performed ROI analyses in major WM tracts previously linked to mathematical abilities in TD children and DD samples (Matejko & Ansari, 2015; Moeller et al., 2015; Peters & De Smedt, 2018). Based on these studies we selected bilateral SLF, bilateral ILF, bilateral IFOF, bilateral ATR, bilateral SCR, bilateral IC, bilateral EC, right CST, body and splenium of the CC and forceps major. We defined the ROIs using the two JHU (Johns Hopkins University) atlases in FSL with a threshold of 25%, yielding the core of each of the tracts. We used the thresholded ROI as a mask in randomise and repeated the permutation analyses (i.e., 10 000 permutations for each ROI) for FA, MD, AD and RD. Again, the TFCE method and FWE correction were applied to determine statistically significant differences between groups.

#### 2.4.3. Tractography Analyses

We aimed to complement our analyses using probabilistic tractography. Probabilistic tractography reconstructs the pathways by taking into consideration the noise and the fiber spreading as uncertainty, dealing with crossing fibers in a voxel and calculating the most likely connections from the fiber orientation distributions (Behrens et al., 2003; Tournier et al., 2011). Probabilistic streamline results can be quantified by generating maps, allowing them to be analyzed and compared more readily (Tournier et al., 2011).

We used the software TRACULA (https://surfer.nmr.mgh.harvard.edu/fswiki/Tracula, (Yendiki et al., 2011) to compare the length of automatically reconstructed major WM pathways between groups. TRACULA is a tool for automatic reconstruction of WM pathways based on global tractography (Jbabdi et al., 2007). TRACULA uses priors from an atlas in combination with the individual cortical parcellation and subcortical segmentation from FreeSurfer. We additionally used the FSL tool PROBTRACKX (Behrens et al., 2007; Behrens et al., 2003) to compute connection probability differences between groups for multiple seed regions. We defined these seed regions based on our previously published task fMRI study in the same cohort and selected the left medial prefrontal cortex, right ventromedial prefrontal cortex/anterior cingulate cortex, left hippocampus, bilateral intra parietal sulci and bilateral occipital cortices (Üstün et al., 2019). The fiber orientation distribution was computed using the FSL tool BEDPOSTX (Behrens et al., 2007; Jbabdi et al., 2012) (with up to two fiber directions per voxel using the ball-and-two-stick model) for both TRACULA and FSL PROBTRACKX.

For TRACULA analysis, we focused on tracts for which we had found significant group differences in the TBSS analyses. We then compared the length of the highest probability path between groups. We also calculated the estimated individual total intracranial volumes (eTIV) from the T1-weighted structural images with FreeSurfer (https://surfer.nmr.mgh.harvard.edu) and used the eTIV as covariate in ANCOVA in SPSS v24 to correct for head size as confounding factor, which could influence the tract length. Additionally, we computed connection probability maps using PROBTRACKX (default settings: 5000 sample tracts per seed voxel, step length 0.5 mm, curvature threshold 0.2, maximum 2000 steps per streamline, volume fraction threshold of subsidiary fiber orientations 0.01) as previously described (Finkl et al., 2020; Jeon et al., 2019). Then, we compared the groups’ connection probability maps in independent samples t-test design with permutation tests (10 000 permutations) in FSL randomise and considered those voxels significant which survived p<0.05 with TFCE method and FWE correction.

#### 2.4.4. Quality Checking for FreeSurfer Parcellation Outputs

Because TRACULA used the FreeSurfer (Fischl, 2012) segmentation and parcellation as prior for the WM tract reconstructions, we checked the quality of the FreeSurfer (v6.0) parcellation and corrected the errors following the software recommendations. We re-ran the FreeSurfer segmentation steps for the whole sample after suitable adjustments.

#### 2.4.5. Correlation Analyses

We tested the relationships between WM connectivity measures and behavioral test scores in the whole sample. For this purpose, we computed Pearson-correlation coefficients using SPSS v24. The behavioral test scores came from mathematical tests (MAT, CPT and CPT-subtests), WISC-R subtests (Verbal, Performance, Arithmetic) and reading evaluations. As WM microstructural connectivity measures, we used DTI derived indices (FA, AD, MD and RD values) from the areas which showed significant group differences in the TBSS and eTIV corrected tract length from the probabilistic tractography analysis using TRACULA.

## 3. RESULTS

### 3.1. Results of TBSS Whole Brain Analyses

We aimed to investigate possible structural connectivity differences in children with DD compared to TD peers. We first compared whole brain FA, MD, RD and AD values as indicators of WM microstructural differences. Voxel-wise whole brain TBSS analyses showed no significant FA, MD, AD or RD differences between groups, neither for Controls>Dyscalculics contrast nor Dyscalculics>Controls contrast (p>0.05, FWE corrected for multiple comparisons with TFCE).

### 3.2. Results of TBSS-ROI Analyses

All reported results here have been found as statistically significant at the p< 0.05 level with FWE correction for multiple comparisons with TFCE in the selected ROIs (Table 2, Fig 1, 2, Fig S2, S3, S4 in Supplementary Material). First, we found lower FA and higher RD, MD and AD values in DD compared to TD children for the left SLF/AF (Table 2, Fig 1). AD and MD values were higher in DD compared to TD for the left ATR, bilateral IC and CC-splenium pathways (Table 2, Fig 2, Fig S2, S3 and S4 in Supplementary Material). In addition to the CC-splenium findings, differences were also found in the body of the CC, showing higher AD values in DD than TD children (Table 2, Fig S2 bottom in Supplementary Material). FA values within the right SCR were lower in DD than in TD (Table 2, Fig S3 bottom in Supplementary Material). Finally, small significant differences were also apparent between groups for left SCR (RD values) and right ATR (AD values). No significant group differences were found for other ROIs for any DTI measures.

**Table 2.** TBSS-ROI Results: White matter microstructure differences between children with DD and TD controls.

| Pathway | Laterality | DTI-measure | Contrast | Voxels | Peak p-value* | Peak MNI Coordinates (mm) |  |  | Figures |
| --- | --- | --- | --- | --- | --- | --- | --- | --- | --- |
|  |  |  |  |  |  | X | Y | Z |  |
| SLF/AF | L | FA | Controls>Dyscalculia | 87 | 0.023 | -38 | -23 | 29 | Figure 1 |
| SLF/AF | L | MD | Dyscalculia>Controls | 256 | 0.016 | -32 | -19 | 31 |  |
| SLF/AF | L | RD | Dyscalculia>Controls | 72 | 0.035 | -34 | -25 | 35 |  |
| SLF/AF | L | AD | Dyscalculia>Controls | 5 | 0.046 | -37 | -14 | 31 |  |
| ATR | L | AD | Dyscalculia>Controls | 195 | 0.003 | -18 | 8 | 7 | Figure 2 |
| ATR | L | MD | Dyscalculia>Controls | 151 | 0.02 | -11 | -3 | 2 |  |
| CC-Splenium | Bilateral | AD | Dyscalculia>Controls | 161 | 0.008 | -1 | -36 | 17 | Figure S2 |
| CC-Splenium | Bilateral | MD | Dyscalculia>Controls | 121 | 0.021 | -7 | -38 | 20 |  |
| CC-Body | Bilateral | AD | Dyscalculia>Controls | 198 | 0.025 | -1 | -12 | 26 | Figure S2 |
| IC | R | AD | Dyscalculia>Controls | 352 | 0.016 | 15 | -7 | 0 | Figure S3 |
| IC | R | MD | Dyscalculia>Controls | 294 | 0.016 | 22 | -7 | 12 |  |
| IC | L | AD | Dyscalculia>Controls | 197 | 0.02 | -10 | -2 | 1 | Figure S4 |
| IC | L | MD | Dyscalculia>Controls | 77 | 0.036 | -19 | 3 | 12 |  |
| SCR | R | FA | Controls>Dyscalculia | 88 | 0.024 | 27 | -11 | 20 | Figure S3 |
| SCR | L | RD | Dyscalculia>Controls | 14 | 0.048 | -25 | -9 | 33 | - |
| ATR | R | AD | Dyscalculia>Controls | 4 | 0.047 | 17 | 7 | 7 | - |

\*: Corrected for multiple comparisons with FWE correction at $p < 0.05$ level with TFCE method. FA: Fractional Anisotropy, AD: Axial Diffusivity, MD: Mean Diffusivity, RD: Radial Diffusivity. SLF/AF: Superior Longitudinal Fasciculus/ Arcuate Fasciculus, ATR: Anterior Thalamic Radiation, CC: Corpus Callosum, IC: Internal Capsule, SCR: Superior Corona Radiata.

**Fig 1.**
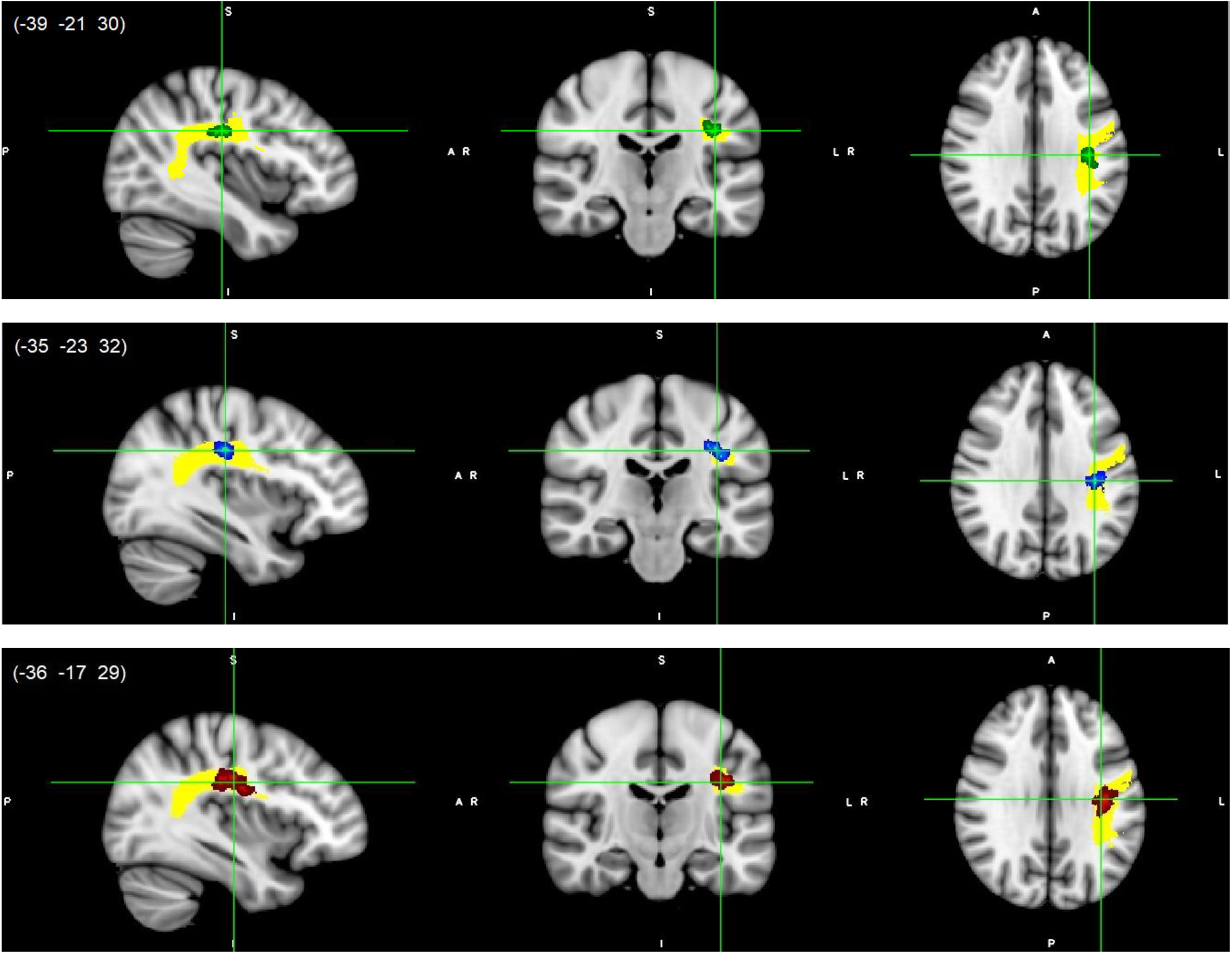
Left SLF white matter microstructure differences. Lower FA (green, top), higher RD (blue, middle) and higher MD (red, bottom) values in DD compared to TD children, p<0.05, FWE corrected with TFCE. Results are thickened with tbss_fill. Yellow region represents selected SLF/AF mask as ROI. SLF/AF, superior longitudinal fasciculus/arcuate fasciculus.

**Fig 2.**
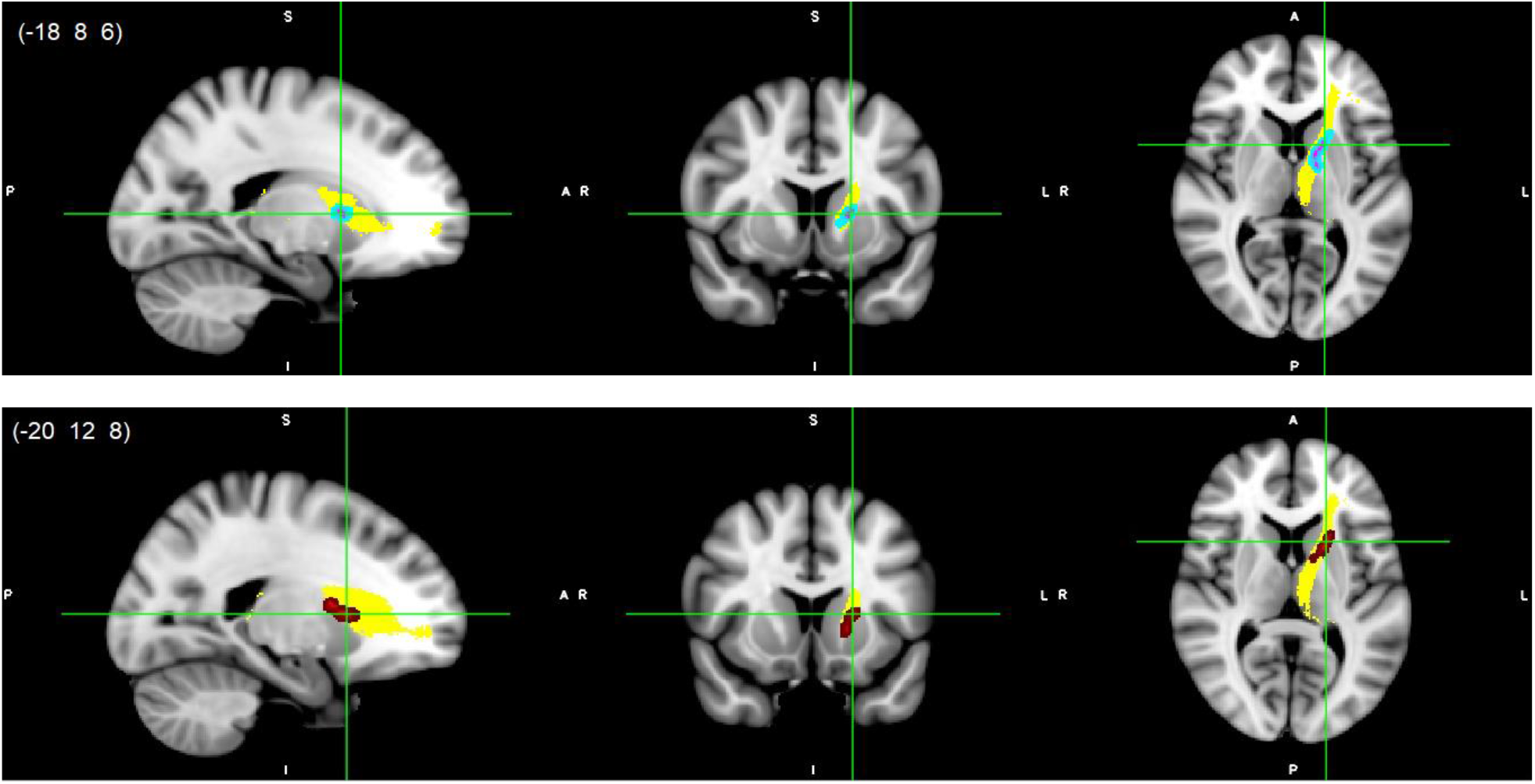
Left ATR white matter microstructure differences. Higher AD (light blue-pink, top) and higher MD (red, bottom) values in DD compared to TD children, p<0.05, FWE corrected with TFCE. Results are thickened with tbss_fill. Yellow region represents selected ATR mask as ROI. ATR, anterior thalamic radiation.

### 3.3. Tractography Results

We performed TRACULA to reconstruct individual pathways using global brain probabilistic tractography using the cortical parcellation from FreeSurfer. We investigated those association and projection fiber tracts showing significant WM microstructure differences between groups in the TBSS-ROI analyses. We compared the fiber length of the left SLF-temporal/AF, left SLF-parietal and left ATR between groups in ANCOVA using individual eTIV as covariate. The left ATR and left SLF-temporal/AF were statistically significantly shorter in DD than TD controls when eTIV was controlled [F (1,23) = 6.26, p = 0.02, partial η^2^ = 0.21 for ATR and F (1,23) = 6.35, p = 0.019, partial η^2^ = 0.22 for SLF-temporal/AF tract]. Reconstructed pathways of the left SLF-temporal/AF and left ATR for each group and tract length differences between groups can be seen in Fig 3. We also depicted the endpoints (i.e. two termination regions) of the reconstructed pathways for each group (see Supplementary Figures S5 and S6). No significant group differences were found in the left SLF-parietal tract: F (1,23) = 1.88, p = 0.18, partial η^2^ = 0.08]. See Supplementary Material Note S3 for means, standard deviations and independent samples t-test statistics of the groups’ length values. Connection probability maps from seed regions obtained via FSL PROBTRACKX did not differ between groups (all p>0.05, FWE-corrected for multiple comparisons with TFCE).

**Fig 3.**
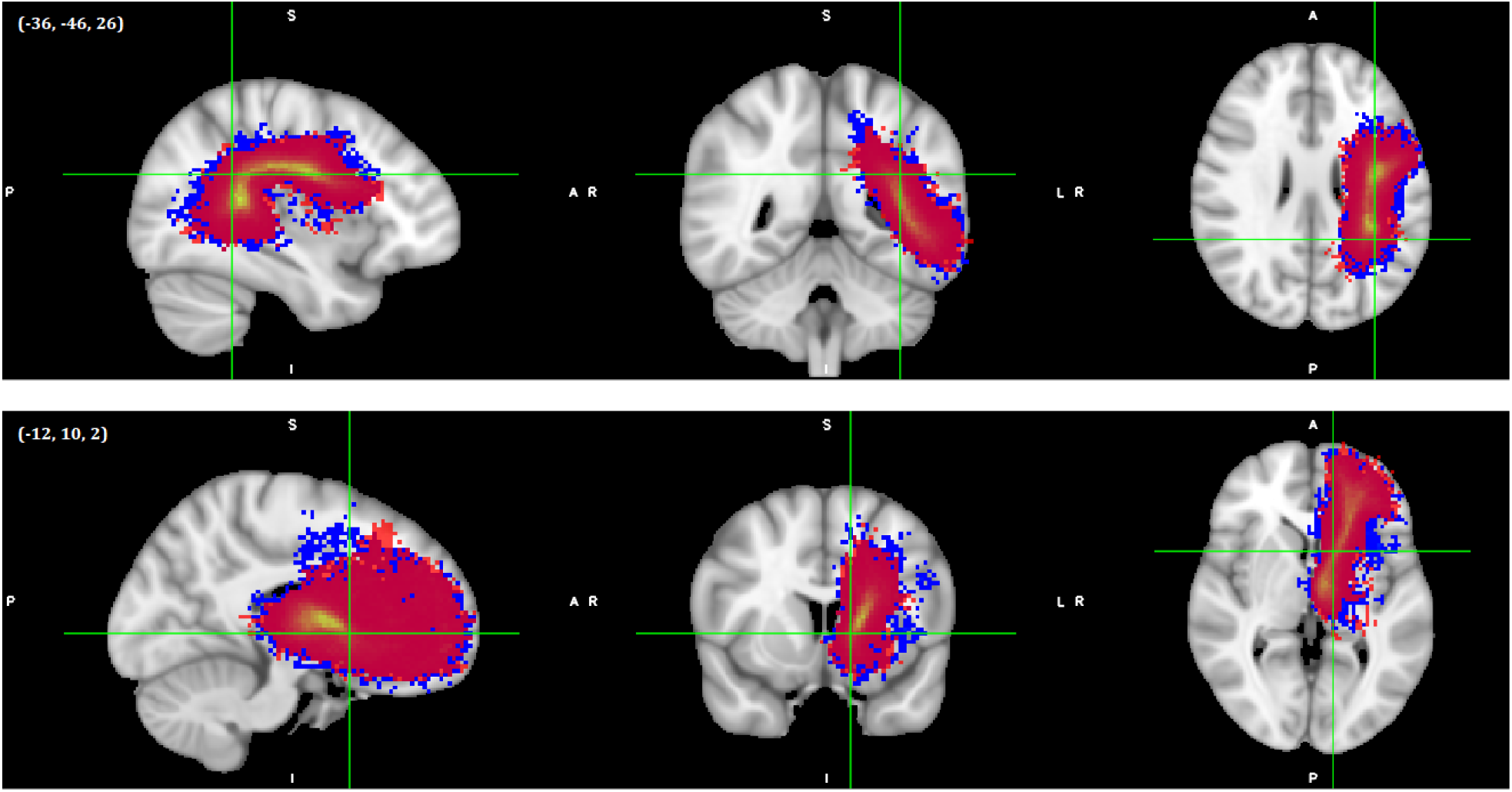
Reconstructed pathways of the DD and TD groups after global tractography. The left SLF-temporal/AF (top) and left ATR (bottom). Red colors represent DD and blue colors represent TD samples. Images shown here are on MNI152 template. In the scope of TRACULA analyses, Freesurfer bbregister was used for registration of diffusion images to T1-weighted images as intra-subject registrations and FSL FLIRT linear registration was used for T1-weighted images to MNI152 template as inter-subject registrations.

### 3.4. Results of the Correlation Analyses

We found significant positive correlations between the mathematical test scores and tract length values of the left SLF-temporal/AF and left ATR tracts. Correlations between math scores, tract length values and DTI derived indices are shown in Fig 4. As shown in Fig 4A, FA values of the left SLF-temporal/AF were positively correlated with the tract length and mathematical test scores, whereas diffusivity values (AD, MD and/or RD) were negatively correlated with the lengths of the reconstructed pathways (see Fig 4B and 4C) and also with the mathematics achievement scores. Other calculation tests and WISC-Arithmetic sub-tests also showed similar correlation patterns (see Supplementary Table S2). Performance IQ and reading scores, on the other hand, were not significantly correlated with any DTI indices or tract length values.

**Fig 4.**
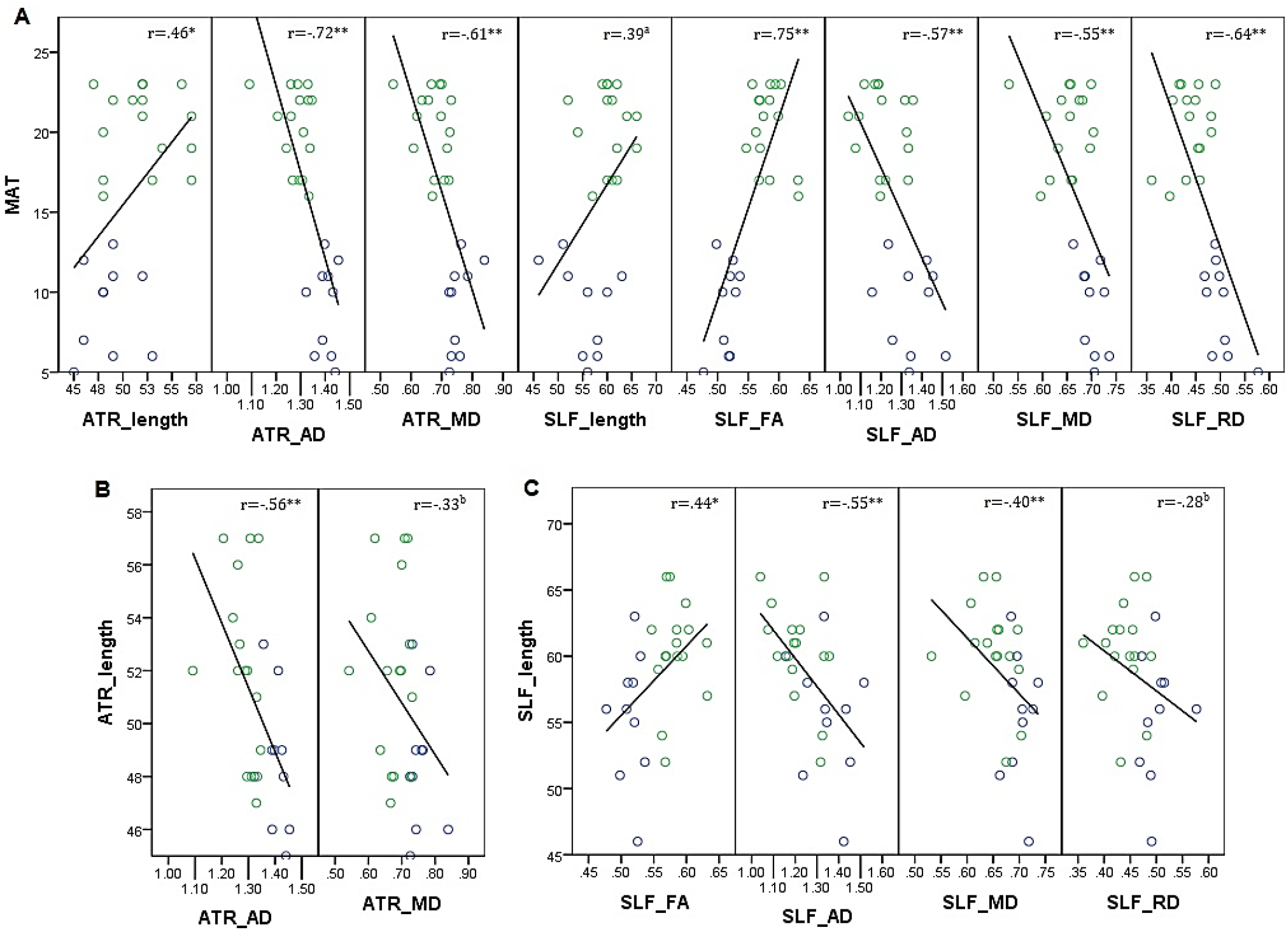
Correlations between white matter coherence (i.e. DTI derived FA, AD, MD, RD values), tract length and math achievement. MAT: Math Achievement Test, ATR: Anterior Thalamic Radiation, SLF: Superior Longitudinal Fasciculus, FA: Fractional Anisotropy, AD: Axial Diffusivity, MD: Mean Diffusivity, RD: Radial Diffusivity, *: p<0.05, **: p<0.01, a: p=0.052, b: p>0.05, blue and green circles represent DD and TD groups, respectively, Unit of the diffusivity values: 10^−3^ mm^2^/s, length measures represent voxel numbers, voxel dimension=1.55mm^3^.

## 4. DISCUSSION

Previous correlational studies in children revealed that numerical abilities relate to WM microstructure of certain association (e.g., bilateral SLF, bilateral ILF, bilateral IFOF), projection (e.g., bilateral ATR, bilateral SCR, bilateral IC, bilateral EC, right CST) and commissural fibers (e.g., body and splenium of the CC and forceps major). We therefore expected structural connectivity alterations in children with DD in these fiber pathways compared to TD peers. Besides WM microstructural properties, for the first time we assessed lengths of the reconstructed pathways and connection probability maps of functionally defined seed regions via probabilistic tractography. In whole brain TBSS and seed-based probabilistic tractography analyses, there were no significant FWE corrected group differences. This might be explained by the small-to-moderate sample size and the resulting low statistical power. However, ROI analyses showed statistically significant reduced WM coherence in the left SLF, left ATR, bilateral SCR, bilateral IC and body and splenium of the CC in the DD group showing consistent lower FA and/or higher diffusivity values. There were no significant WM microstructural differences in the bilateral IFOF, bilateral ILF, right SLF, right CST, bilateral external capsules and forceps major ROIs, which is in contrast to previous studies in children (Hu et al., 2011; Lebel et al., 2010; Li, Hu, et al., 2013; Li, Wang, et al., 2013; Navas-Sanchez et al., 2014; Rykhlevskaia et al., 2009; Tsang et al., 2009).

We provided the first evidence that DD children had statistically significant shorter ATR and shorter SLF-temporal/AF in the left hemisphere when controlling for intracranial volumes. Previous studies investigating the development of the pathways (i.e. the limbic and cerebellar pathways) from childhood to young adulthood showed that age had a positive correlation with tract length (Re et al., 2017; Yu et al., 2014). Thus, as the brain grows, maturation and myelination rates of the axons increase during normal development, and the tracts get longer with age. On the other hand, in normal aging studies that included autopsied brains from young adulthood to older ages, the total tract lengths were found to decrease with age as the number of the thinner myelinated fibers decreased (Marner et al., 2003; Tang et al., 1997). Likewise, *in vivo* aging studies with tractography showed shorter fiber bundles in healthy older samples (Baker et al., 2014; Bolzenius et al., 2013; Salminen et al., 2013). Several studies also indicated that shorter tracts were associated with neuropsychiatric diseases (Buchsbaum et al., 2006; Torgerson et al., 2013). In those studies, shorter pathways were mirrored to be related to a loss/deficit of myelinated fibers, a reduction in fiber density/number of fibers, a miswiring problem related to less topographically developed fiber tracts or a change in the underlying architecture of the pathway. Altered fiber architecture might result in/from “more poorly organized fibers, fanning out to the cortex irregularly with less topographic precision and perhaps earlier in their course” (Buchsbaum et al., 2006). Shorter pathways in DD, therefore, might be interpreted as weaker communication between cortex areas due to altered/delayed maturation/myelination of the axons or a lack of important longer fibers, or interpreted as altered communication caused by different trajectories between the brain regions. Moreover, lower WM coherence [i.e., lower FA and/or higher diffusivity values] in our DD sample seems to support these interpretations.

Finally, we performed correlation analyses between neuropsychological test scores and WM connectivity measures (i.e. DTI derived indices and tract length measures) to explore the link between imaging biomarkers and behavior in dyscalculia. Some aging and clinical studies found fiber length to be positively correlated with cognitive functions, whereas they were negatively correlated with age or disease condition (Baker et al., 2017; Behrman-Lay et al., 2015; Heaps-Woodruff et al., 2018; Salminen et al., 2013). In our study, we revealed that tract lengths of the left SLF/AF and left ATR were positively correlated with mathematical abilities. Furthermore, the lengths were positively correlated with the DTI derived FA measures and negatively correlated with the diffusivity values. Similar relationships were not present for reading or performance IQ scores, which is in line with the *pure* DD definition. It is important to emphasize that, to our knowledge, there has been no developmental study investigating the relationships of *in vivo* or *ex vivo* fiber lengths and behavior.

There were no tract length differences for the left SLF-parietal tract. In the following, we discuss the results of our regional analyses targeting major fiber tracts related to numerical disabilities.

### 4.1. Association Fibers

We found lower FA, higher RD and higher MD values (also higher AD but only in a small cluster) exclusively in the left SLF/AF in children with DD compared to TD peers. These differences were in the white matter of the parietal lobe adjacent to the central sulcus. Moreover, we found shorter left SLF/AF tract length in DD than in TD children. This might mean that some of the pathways of the left SLF/AF in DD are lacking or cannot reach to the cortex areas and/or they might have a different trajectory from the pathways in the TD group. When we examine the reconstructed left SLF/AF between groups (see Fig 3), the tracts in the TD group seem to have a different trajectory that is more dorsal, lateral and superior than the DD group. Despite no considerable differences, the frontal and temporal endpoints, respectively, appear to be more superior and more dorsal-lateral-posterior in our TD group compared to DD (see Supplementary Fig S5). More dorsal-lateral-superior tracts of the TD group also resemble the longer pathways and that might mean that the TD group has much longer tracts, whereas the DD group has mostly shorter tracts and lacks longer ones. This was also supported by the finding that the minimum and especially maximum tract length values of the TD group were greater than those of the DD group (see Notes S4), although not significant. In line with this, we also showed that shorter SLF/AF tracts are associated with lower math achievement and lower WM coherence in the whole sample.

Our findings are consistent with the results of the previous two studies which investigated WM in DD showing lower FA values in bilateral SLF (Kucian et al., 2014; Rykhlevskaia et al., 2009). Kucian et al. (2014) showed lower FA in the bilateral SLF (FWE corrected and mostly in left) and higher RD (and also lower AD in a small cluster) in the left SLF (uncorrected results) in the whole brain as well as in the ROI analyses in 10-year-old children. Rykhlevskaia et al. (2009), however, found lower FA values especially in the temporo-parietal WM regions (mostly in right) including bilateral SLF in 7–9-year-old children.

In TD children, WM microstructural properties in the SLF were also found to be related to mathematical performance. In 10–15-year-old children, Tsang et al. (2009) showed FA values in bilateral SLF, and left fronto-parietal connections of the SLF were positively correlated with arithmetic scores in TBSS and deterministic tractography, respectively. Li, Hu, et al. (2013) used fiber tracking with the left IPS after VBM analysis and showed that FA values in the left SLF were positively correlated with arithmetic performance in 10-year-old children. Using whole brain tractography, Van Beek et al. (2014) found that higher FA in the left arcuate fasciculus specifically predicted better addition and multiplication performance in 12-year-old children. Jolles et al. (2016) examined learning-related changes in 7.7–9.1-year-old children and found increased FA changes in the fronto-temporal part of the left SLF.

Impaired arithmetic competencies in other clinical conditions in children and adolescents are also related to reduced FA in the left SLF (Barnea-Goraly et al., 2005; Lebel et al., 2010; Till et al., 2011). Math-gifted adolescents had higher FA in the bilateral SLF compared to TD controls (Navas-Sanchez et al., 2014). Additionally, arithmetic performance of young adults was correlated positively with FA or negatively with RD also in the left SLF (Klein et al., 2013; Matejko et al., 2013).

The SLF connects frontal regions to parietal-temporal and occipital regions (Conner et al., 2018; Makris et al., 2005; Schmahmann et al., 2008). The fronto-parieto-temporal part of the SLF is known as the AF and is involved in the dorsal stream of the language network in the brains of adults and children (Anwander et al., 2007; Brauer et al., 2011; Catani et al., 2005; Eichert et al., 2019; Wang et al., 2016). The dorsal language network is related to the phonological aspects and verbal working memory processes of language (Friederici & Gierhan, 2013; Hickok & Poeppel, 2004). In addition to bilateral IPS and bilateral superior parietal lobules for number processing, concordant with the dorsal language network, verbal numerical processing has been attributed to the left perisylvian language regions (mainly to left angular gyrus) (Dehaene, 2011; Dehaene et al., 2003). Task-based fMRI meta-analyses have shown the fronto-parieto-temporal network is involved in the brain’s numerical functions (Arsalidou et al., 2018; Arsalidou & Taylor, 2011; Houde et al., 2010; Kaufmann et al., 2011; Sokolowski et al., 2017). Present results showing both reduced WM coherence and shorter length of the left SLF/AF therefore might be associated with verbal number processing problems in DD.

### 4.2. Projection Fibers

We found lower FA in the right SCR, higher RD in the left SCR (in a relatively small cluster) and higher AD and higher MD in the bilateral IC in DD compared to TD peers. In line with our findings, Tsang et al. (2009) found that FA values in the corona radiata and IC were positively correlated with arithmetical performance in TD children. Similarly, Van Eimeren et al. (2008) showed that FA in the left SCR positively correlated with numerical operations and mathematical reasoning in 7–9-year-old TD children. Hu et al. (2011) found reduced RD mostly in the right IC and right corona radiata after math training in TD children. Lebel et al. (2010) found lower FA in bilateral IC (also in the left CST) in children with dyscalculia in another clinical sample. IC and SCR extend throughout the sensory-motor pathways from/to high level cognitive brain areas (FitzGerald et al., 2012). Some studies show that sensory-motor pathways could be related to finger counting (Noel, 2005; Sato et al., 2007). Therefore, differences in IC and SCR might be related to deficiencies in DD regarding use of counting strategies in arithmetical problem-solving.

Our findings showed that AD and MD in the left ATR (also a small cluster in the right ATR) were larger in the DD group. Additionally, we found a shorter left ATR in children with DD. When we compare the reconstructed left ATR pathways between groups (see Fig 3), the TD group seems to have more tracts connecting the more posterior, lateral and inferior parts of the prefrontal cortex. Namely, the tracts in the TD group connect more extended areas including more branches to the prefrontal regions than the tracts in the DD group. This might also be caused by the different developmental stages of the prefrontal cortex, similar to the growth of the cingulum tract from birth through young adulthood (Yu et al., 2014). The prefrontal endpoints of the TD group also appear to be more inferior and lateral in the TD compared to the DD group (see center of gravity coordinates in Supplementary Fig S5). This might mean that the TD group has much longer tracts reaching to the prefrontal cortex compared to the DD group. The maximum tract length values were also greater in the TD group (see Notes S4), although not significant. Furthermore, we showed that the shorter ATR tracts were correlated with lower math achievement and lower WM coherence in the whole sample.

Similar to our findings, Rykhlevskaia et al. (2009) found lower FA in the bilateral ATR in their DD sample. Moreover, Navas-Sanchez et al. (2014) showed higher FA values in bilateral thalamic radiations (also in bilateral IC) in math-gifted adolescents. The ATR has reciprocal connections from/to the anterior nuclei of the thalamus, which is a conjunction hub for the connections between hippocampus and mammillary bodies, to/from the (pre)frontal as well as anterior cingulate cortices (Behrens et al., 2003; Child & Benarroch, 2013). This tract has been shown to be related to learning, episodic and spatial memory and executive functions of the brain (Aggleton et al., 2010; Dillingham et al., 2015; Jankowski et al., 2013; Vanderwerf et al., 2003). Findings in our study showing both reduced WM coherence and shorter length in the left ATR might explain why learning, memorizing and processing numbers are problematic in DD. This is supported by our previously published task based fMRI study in which bilateral prefrontal regions were hyper-activated in DD in a symbolic and non-symbolic number comparison task, and specifically the left hippocampus was hyper-activated in DD in the symbolic number comparison task (Üstün et al., 2019). These hyper-activations, interpreted as compensatory mechanisms in DD, in the left hippocampal and prefrontal regions might be associated with the left ATR structural connectivity differences in this study.

### 4.3. Commissural Fibers

We found higher AD and RD in the body and splenium parts of the CC in DD compared to TD children. Consistent with our findings, Rykhlevskaia et al. (2009) found lower FA in the splenium connections of their DD sample. Dyscalculic children and adolescents with other clinical diseases also showed positive correlations between FA in the CC and math/arithmetic scores (Lebel et al., 2010; Till et al., 2011). In TD children, Tsang et al. (2009) showed that FA values in the splenium were positively correlated with approximate arithmetic performance. After math training, TD children had higher FA values in the splenium and body of the CC (Hu et al., 2011). Moreover, Navas-Sanchez et al. (2014) revealed that math-gifted children had higher FA in the splenium of the CC (also in the forceps minor) compared to TD children. Cantlon et al. (2011) investigated interhemispheric IPS connectivity with deterministic tractography, revealing that FA in the left isthmus (part of the body) of the CC predicted number magnitude comparison performance in 6-year-old children. The splenium of the CC primarily connects temporo-parietal and also occipital lobes; the body of the CC connects primary and secondary sensory-motor cortices (Aboitiz et al., 1992; Gazzaniga, 2000; Knyazeva, 2013). Differences in bilateral sensory-motor connectivity and the reduced fronto-parieto-temporal connectivity in our study might contribute to a reduced interhemispheric communication in DD in the body and splenium parts, respectively.

## 5. LIMITATIONS

The main limitation of this study is the small sample size. This was because we could only re-assess the children after two years instead of one year because of uncontrollable external reasons during the time of the study. This delay made it impossible to include all selected children because some had already finished primary school and were not reachable anymore due to the data protection reasons. Another limitation of the study is that we did not formally assess socioeconomic status (SES) of the children, which might be an important confounder regarding math development but also measures of brain structure (Demir-Lira et al., 2016; Noble et al., 2012; Noble et al., 2005; Rosen et al., 2018).

For TBSS-ROI analyses, we defined 25% of the maximum intensity values of each tract as thresholds in the FSL JHU WM tractography atlas by inspecting them carefully. Yet, every voxel in the mean FA skeleton might not have totally overlapped with the ROI mask. Additionally, body and splenium parts of the CC, IC and EC ROIs were chosen from FSL JHU ICBM81 WM atlas. For TRACULA analyses, we used only the SLF and the ATR as ROIs because TRACULA has no implementation for the other pathways which differed between groups in TBSS-ROI analyses.

Finally, even though commonly used, shortcomings of DTI include potential artefacts due to crossing fibers within a voxel as well as partial volume effects (Jones et al., 2013; Tournier et al., 2011). It must be emphasized that DTI constructs a model of “brain fibers” which is constantly being improved with advancing technology. However, due to the limited spatial resolution of MRI and the aforementioned issues, DTI still cannot be equated with the “true” biological basis.

## 6. CONCLUSIONS

We studied structural connectivity underlying *pure* DD in children. One main finding of our study—reduced WM coherence and shorter length of the left superior-longitudinal/arcuate-fasciculus—might be related to deficiencies in verbal aspects of number processing in DD. The other major finding—reduced WM coherence and shorter length of the left anterior thalamic radiation—might be attributed to insufficiencies in encoding, retrieving and working with arithmetical facts in children with DD. Left lateralized differences in these pathways might be related to hemispheric dominance, which was proposed to be pronounced in arithmetic processing during development (Artemenko et al., 2020). In this neurodevelopmental study, we revealed first evidence that lower math performance corresponded with lower WM coherence and shorter pathways. However, further studies are necessary to investigate DD in children with larger data sets, using longitudinal and interventional designs to confirm these findings.

## Acknowledgements

We thank all children, parents and teachers for voluntary participation in this study. We also thank the officers and administration staff of the Ankara Provincial Directorate for National Education.

## Ethics approval

All procedures performed in the study involving human participants were in accordance with the principles of the Declaration of Helsinki. This study was approved by the Ethical Committee of the Faculty of Medicine at Ankara University.

## Consent to participate

Written informed consents were obtained from the parents and all individual participants included in the study.

## Availability of data and material

Applicable after publication of the manuscript.

## Code availability

Applicable with email to the corresponding author.

## Authors’ Contributions

Writing - original draft preparation: [Nazife Ayyıldız]; Writing - review and editing, Data curation, Investigation, Methodology: [Nazife Ayyıldız], [Frauke Beyer], [Sertaç Üstün], [Emre H. Kale], [Öykü Mançe Çalışır], [Pınar Uran], [Özgür Öner], [Sinan Olkun], [Alfred Anwander], [A. Veronica Witte], [Arno Villringer], [Metehan Çiçek]; Formal analysis, Visualization, Software: [Nazife Ayyıldız], [Frauke Beyer], [Alfred Anwander], [A. Veronica Witte]; Conceptualization: [Metehan Çiçek], [Sinan Olkun], [Özgür Öner]; Resources, Supervision, Funding Acquisition: [Metehan Çiçek], [Arno Villringer]; Project administration: [Metehan Çiçek].

