## Supplementary Tables, Notes and Figures for "Structural brain connectivity in children with developmental dyscalculia"

### Supplementary Tables and Notes

**Table S1**. Eddy quality assessment parameters before and after parameter adaptation and exclusion of the bad data.

| | **QA* values of the whole sample (N=12/DD: N=16/Control)** | | **QA values after exclusion of two data sets and changing outlier SD (N=10/DD: N=16/Control)** | |
|---|---|---|---|---|
| **Parameters** | Minimum | Maximum | Minimum | Maximum |
| Signal-to-Noise (SNR) (for b0 images) | 13.11 | 25.88 | 13.11 | 25.88 |

| | | | | |
|---|---|---|---|---|
| Contrast-to-Noise (CNR) (for DWI images) | **0.42** | 1.06 | **0.58** | 1.06 |
| Absolute Motion (mm) | 0.18 | **4.18** | 0.18 | **1.91** |
| Relative Motion (mm) | 0.08 | **2.28** | 0.08 | **1.04** |
| Total Outliers Percentage (%) | 0 | **4.89** | 0 | **2.33** |
| (Slice-to-Volume) Translation (x, y, z) (mm) | (0) 0 | **(<1.5) <3** | (0) 0 | **(0.7) <1.5** |
| Rotation (x y, z) (degree) | (0) 0 | **(<1.5) <2** | (0) 0 | **(0.5) <1.5** |
| Eddy Current Linear Terms | 0 | **0.25** | 0 | **0.04** |

*: Quality Assurance, FSL Eddy QUAD and SQUAD parameter results

**Table S1 (Note S1).** As a conclusion for the processes of the pre-processing steps, we finally included 10 DD and 16 TD children's DTI data. We used the default parameter for the detection of outliers defined by the slice intensity being 4 standard deviations (SDs) below the mean for outlier detection and replacement parameter except for one subject whose SD threshold was increased to 5 to keep all slices without visible artefacts. Thus, maximum total outlier percentage in the data was 2.33 % in all slices in all volumes for any subject. Minimum CNR value was 0.58 and maximum absolute and relative motion measures were 1.91mm and 1.04mm, respectively. The maximum relative motion in the data was less than the voxel size (i.e., 1.55mm).

**Table S1 (Note S2).** Despite no significant differences between groups before excluding the two outlying subjects [for absolute motions t (26) = -1.37, p= 0. 141, means±SD 0.7±0.45mm for TD and 1.36±1.41mm for DD; for relative motions t (26) = -1.57, p= 0. 195, means±SD 0.23±0.23mm for TD and 0.49±0.64mm for DD], after excluding the two outlying subjects the t-scores and variance differences between DD and TD decreased considerably [for absolute motions t (24) = -0.49, p= 0.634, means±SD 0.7±0.45mm for TD and 0.81±0.63mm for DD; for relative motions t (24) = -0.23, p= 0.822, means±SD 0.23±0.23mm for TD and 0.24±0.15mm for DD].

**Note S3:** The left ATR and left SLF-temporal were statistically significantly shorter in DD than TD controls in independent t-test designs ([t (24) = 2.81, p = 0.01, mean±SD 52.1±3.5 for TD and 48.5±2.6 for DD] and [t (24) = 2.88, p =0.008, mean±SD 60.4±3.7 for TD and 55.5±4.9 for DD], respectively, Bonferroni corrected p = 0.017). There were no significant group differences in the left SLF-parietal tract [t (24) = -0.73, p = 0.47, mean±SD 43.63±4 for TD and 44.8±4.05 for DD].

**Note S4.** Minimum and maximum tract length values (i.e. number of voxels, 1voxel=1.55mm$^3$) of the groups.

The left SLF/AF tract length values of the TD group:

Minimum length range= [35-46], mean±SD=38.63±2.85 and maximum length range= [85-122], mean±SD=102±10.42.

The left SLF/AF tract length values of the DD group:

Minimum length range= [33-39], mean±SD=36.7±2.1 and maximum length range= [83-100], mean±SD=94.2±5.55.

ANCOVA results:

No significant group differences for the minimum and maximum lengths of the left SLF/AF tract controlling the eTIV, respectively, $F(1,23) = 1.84$, $p = 0.188$, partial $\eta^2 = 0.074$ and $F(1,23) = 2.98$, $p = 0.098$, partial $\eta^2 = 1.12$.

The left ATR tract length values of the TD group:

Minimum length range= [26-38], mean±SD=30.8±3.1 and maximum length range= [77-119], mean±SD=93.8±13.18.

The left ATR tract length values of the DD group:

Minimum length range= [23-33], mean±SD=29.6±3.1 and maximum length range= [77-108], mean±SD=92.2±10.03.

ANCOVA results:

No significant group differences for the minimum and maximum lengths of the left ATR tract controlling the eTIV, respectively, $F(1,23) = 0.35$, $p = 0.554$, partial $\eta^2 = 0.015$ and $F(1,23) = 0.006$, $p = 0.94$, partial $\eta^2 = 0.00025$.

### Supplementary Figures

**Fig S1**

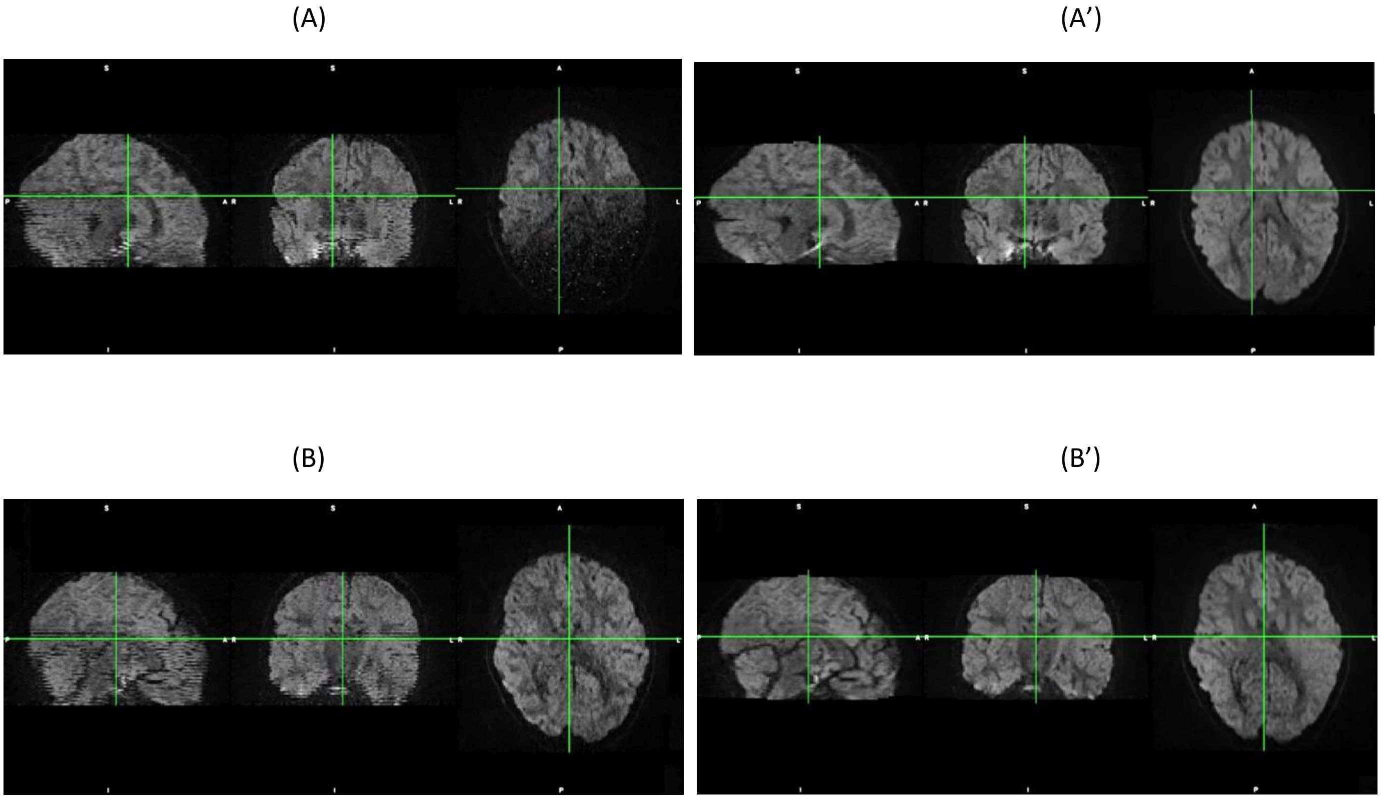


**Fig S1** (A and B) show images with slices showing a signal drop-out (partly black slice in A) and zig-zag patterns occurred (A and especially in B) because of excessive movements between slices and within volumes. After eddy-correction with outlier replacement and slice-to-volume corrections, the signal was reconstructed resulting in no signal loss in A’ and no zig-zag patterns in A’ and B’.

**Fig S2**

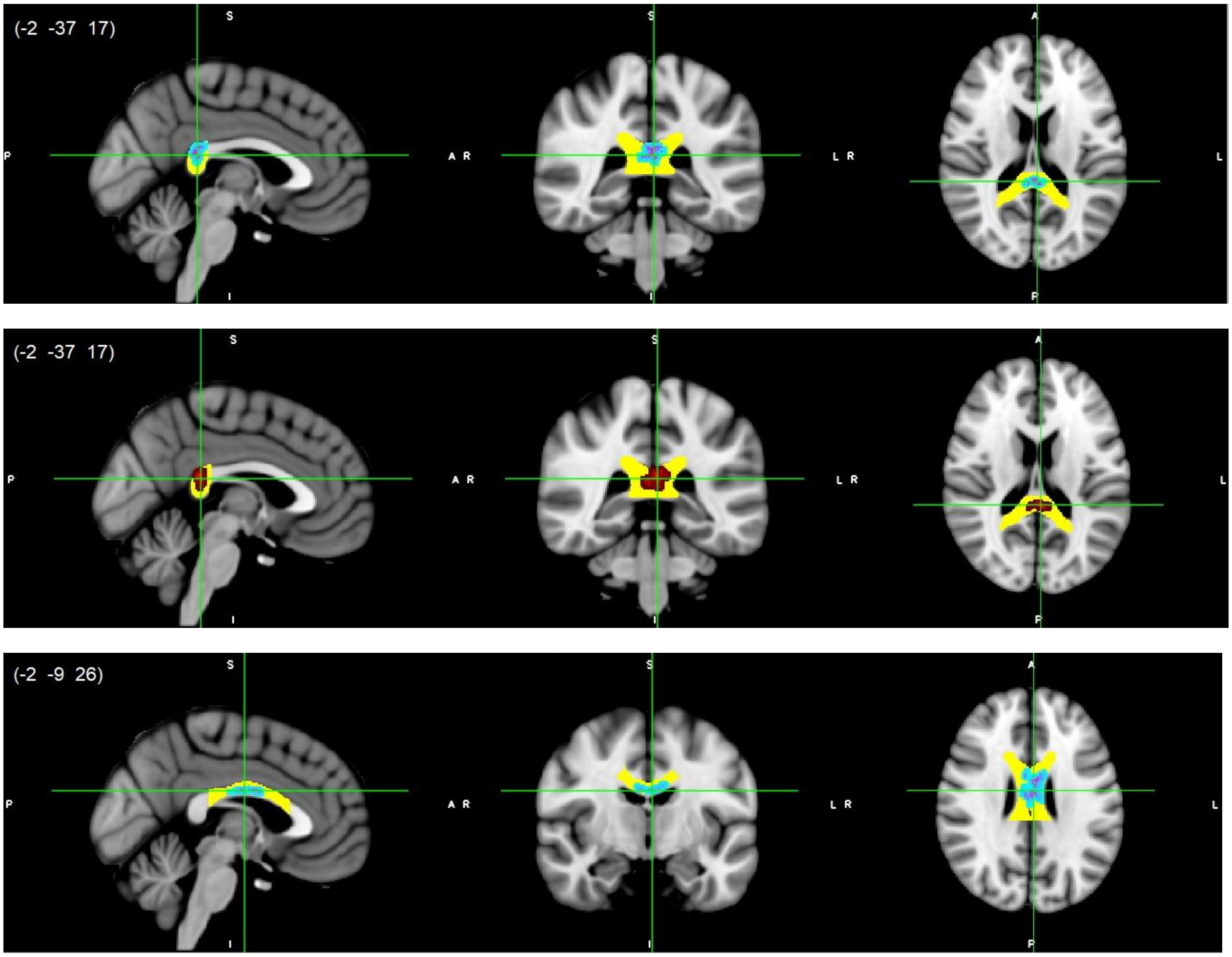


**Fig S2** CC splenium (top/middle) and body (bottom) white matter microstructure differences. Higher AD (light blue-pink, top and bottom) and higher MD (red, middle) values in DD compared to normally developing children, $p<0.05$, FWE corrected with TFCE. Results are thickened with tbss_fill. Yellow regions represent selected CC masks as ROIs. CC, corpus callosum.

**Fig S3**

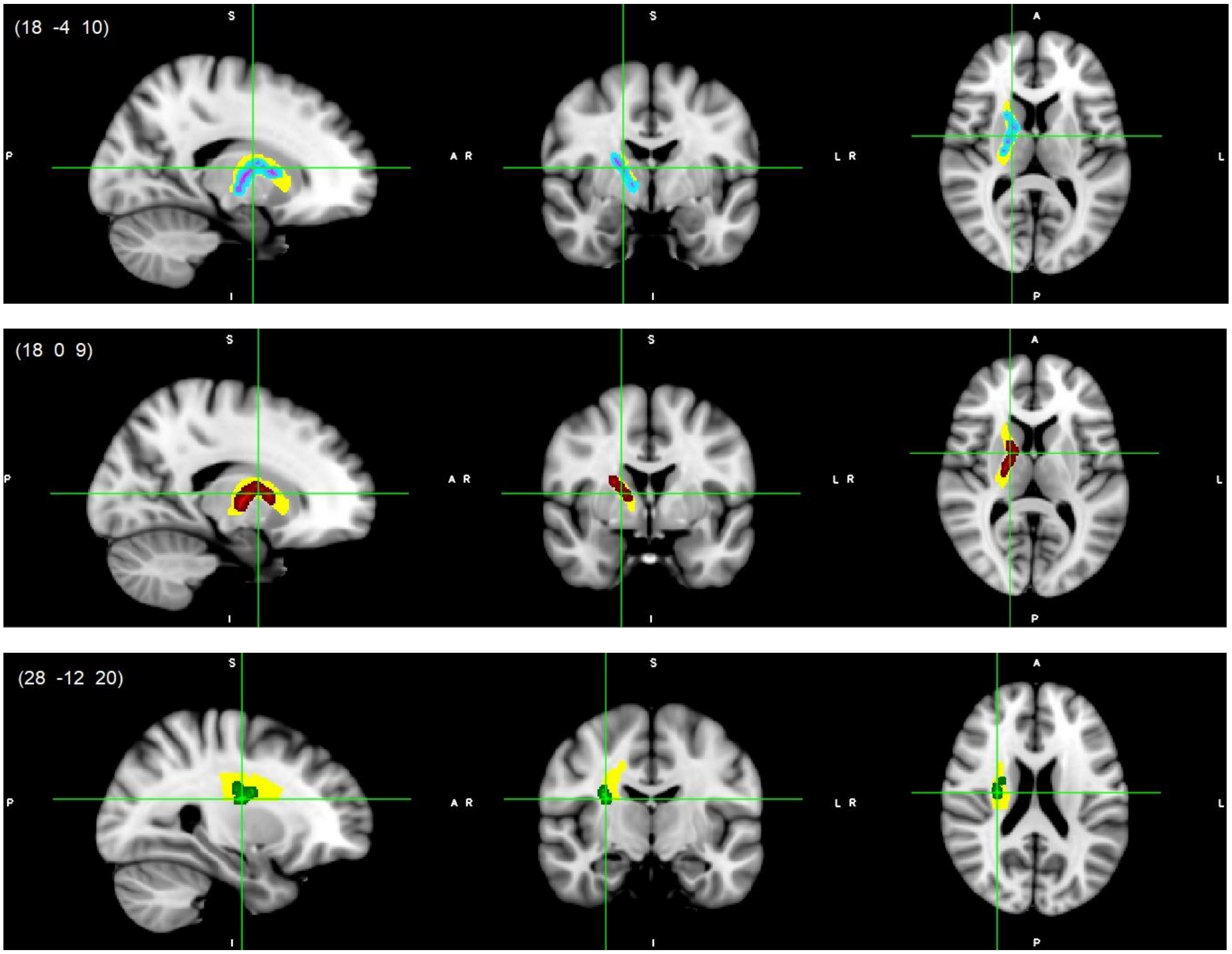


**Fig S3** Right IC (top/middle) and right SCR (bottom) white matter microstructure differences. Higher AD (light blue-pink, top), higher MD (red, middle) and lower FA (green, bottom) values in DD compared to normally developing children, $p<0.05$, FWE corrected with TFCE. Results are thickened with tbss_fill. Yellow regions represent selected IC and SCR masks as ROIs. IC, Internal Capsule; SCR, Superior Corona Radiate.

**Fig S4**

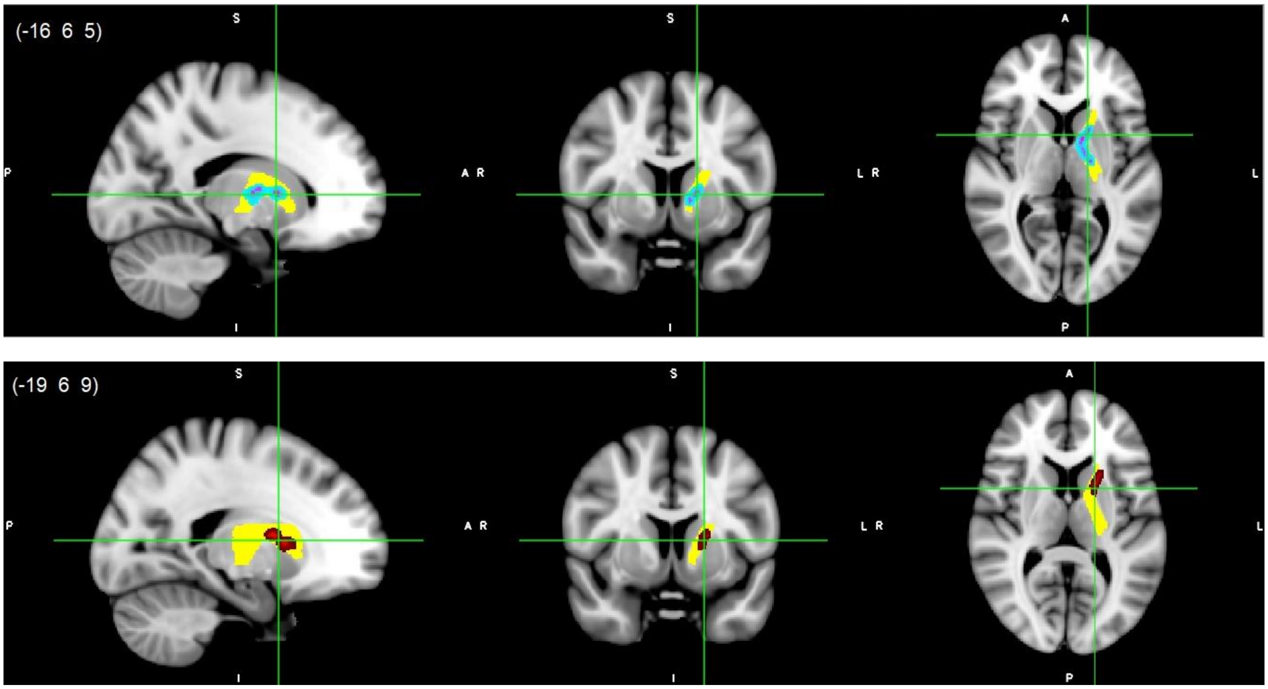


**Fig S4** Left IC white matter microstructure differences. Higher AD (light blue-pink, top) and higher MD (red, bottom) values in DD compared to normally developing children, p<0.05, FWE corrected with TFCE. Results are thickened with tbss_fill. Yellow region represents selected IC mask as ROI. IC, Internal Capsule.

### Fig S5

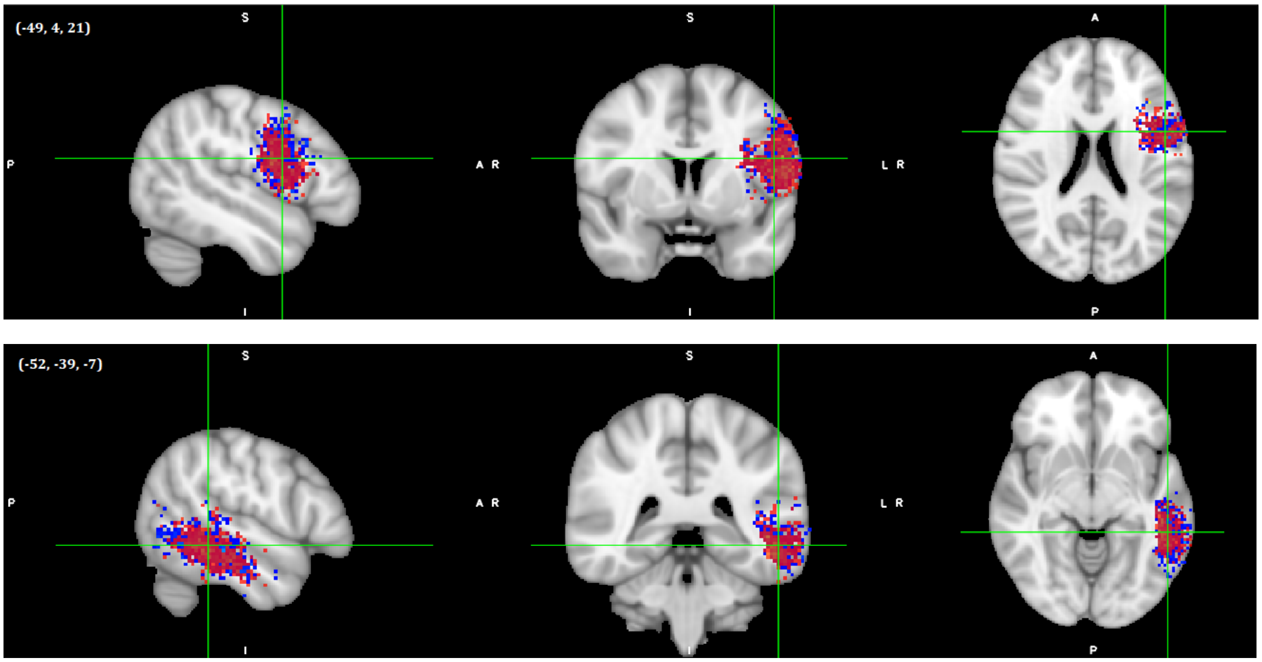


**Fig S5** Endpoints for the left SLF-temporal/AF pathway reconstruction in global tractography analyses. Frontal endpoint (top) and temporal endpoint (bottom). Red colors represent DD and blue colors represent TD samples. Images shown here are on MNI152 template. Center of gravity coordinates of the frontal endpoints were (-49.1, 3.8, 21.3) and (-49.2, 4.4, 19.6) for TD and DD groups, respectively. Center of gravity coordinates of the temporal endpoints were (-52.1, -38.9, -7.1) and (-52.2, -37.2, -8.2) for TD and DD groups, respectively.

**Fig S6**

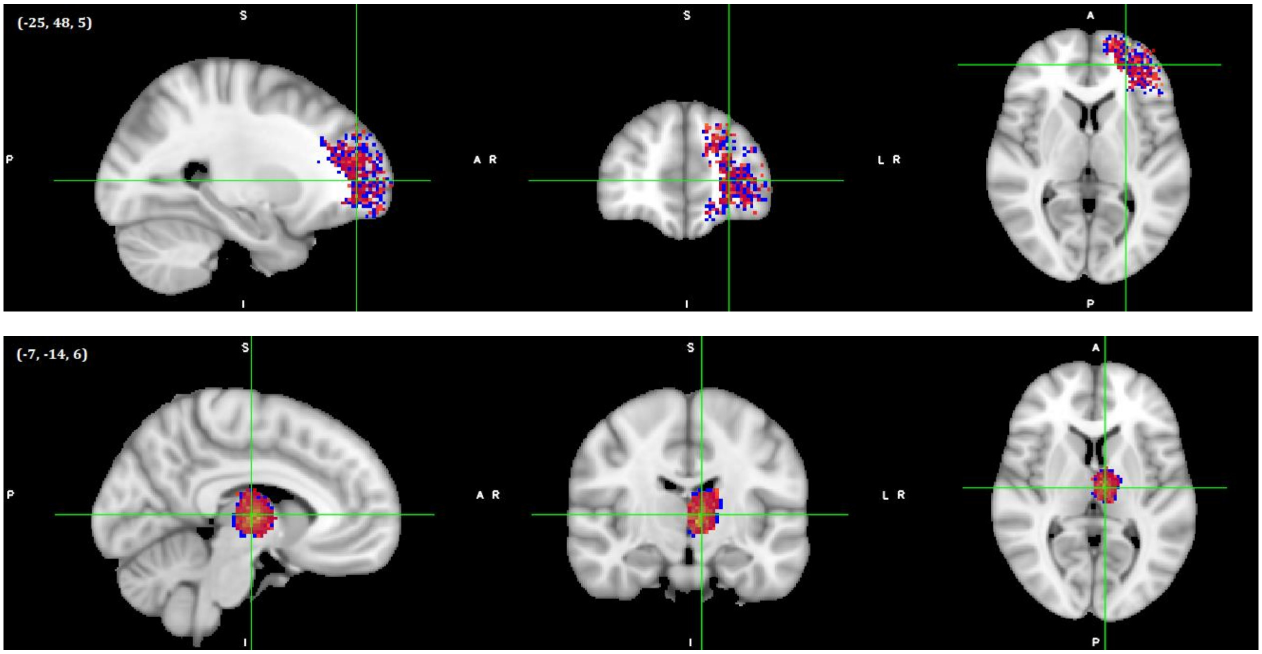


**Fig S6** Endpoints for the left ATR pathway reconstruction in global tractography analyses. Prefrontal endpoint (top) and anterior thalamus endpoint (bottom). Red colors represent DD and blue colors represent TD samples. Images shown here are on MNI152 template. Center of gravity coordinates of the prefrontal endpoints were (-25.2, 47.9, 5) and (-25.9, 47.5, 8.3) for TD and DD groups, respectively. Center of gravity coordinates of the thalamus endpoints were (-6.8, -13.8, 6) and (-6.8, -13.8, 5.9) for TD and DD groups, respectively.
